# H3.3K4M destabilizes enhancer epigenomic writers MLL3/4 and impairs adipose tissue development

**DOI:** 10.1101/301986

**Authors:** Younghoon Jang, Chaochen Wang, Aaron Broun, Young-Kwon Park, Lenan Zhuang, Ji-Eun Lee, Eugene Froimchuk, Chengyu Liu, Kai Ge

**Affiliations:** Adipocyte Biology and Gene Regulation Section, Laboratory of Endocrinology and Receptor Biology, National Institute of Diabetes and Digestive and Kidney Diseases, National Institutes of Health, Bethesda, MD 20892; Transgenic Core, National Heart, Lung, and Blood Institute, NIH, Bethesda, MD 20892

## Abstract

Histone H3K4 mono-methyltransferases MLL3 and MLL4 (MLL3/4) are required for enhancer activation during cell differentiation, though the mechanism is incompletely understood. To address MLL3/4 enzymatic activity in enhancer regulation, we have generated two mouse lines: one expressing H3.3K4M, a lysine-4-to-methionine (K4M) mutation of histone H3.3 that inhibits H3K4 methylation, and the other carrying conditional double knockout of MLL3/4 enzymatic SET domains. Expression of H3.3K4M in lineage-specific precursor cells depletes H3K4 methylation and prevents adipogenesis and adipose tissue development. Mechanistically, H3.3K4M prevents enhancer activation in adipogenesis by destabilizing MLL3/4 proteins but not other Set1-like H3K4 methyltransferases. Notably, deletion of the enzymatic SET domain of MLL3/4 in lineage-specific precursor cells mimics H3.3K4M expression and prevents adipose tissue development. Interestingly, destabilization of MLL3/4 by H3.3K4M in adipocytes does not affect adipose tissue maintenance and function. Together, our findings indicate that H3.3K4M destabilizes enhancer epigenomic writers MLL3/4 and impairs adipose tissue development.

## Introduction

During cell differentiation, transcriptional enhancers are bound by lineage-determining transcription factors (LDTFs) and play a key role in regulating cell type-specific gene expression. Cell type-specific enhancers are marked by specific epigenomic features ^1^. Histone 3 lysine 4 (H3K4) mono-methylation (H3K4me1) is the predominant mark of a primed enhancer state. Histone 3 lysine 27 acetylation (H3K27ac) by H3K27 acetyltransferases CBP/p300 further follows H3K4me1 to mark an active enhancer state ^2^. There are six mammalian Set1-like H3K4 methyltransferases, each containing a catalytic SET domain that enables the deposition of methyl marks on H3K4: MLL1 (or KMT2A), MLL2 (or KMT2B), MLL3 (or KMT2C), MLL4 (or KMT2D), SET1A (or KMT2F), and SET1B (or KMT2G) ^3^. Among them, MLL4 is a major mammalian H3K4 mono-methyltransferase with partial functional redundancy with MLL3. MLL3 and MLL4 (MLL3/4) are required for CBP/p300-mediated enhancer activation in cell differentiation and cell fate transition ^4-6^. Deletion of *Mll3/4* genes depletes H3K4me1 in cells and prevents the enrichment of CBP/p300-mediated H3K27ac, epigenome reader BRD4, Mediator coactivator complex, and RNA Polymerase II on enhancers. Consequently, *Mll3/4* deletion prevents enhancer RNA production, cell type-specific gene induction and cell differentiation ^7^. However, the role of H3K4me1 in enhancer regulation, cell differentiation and function *in vivo* is poorly understood.

Adipogenesis and adipose tissue are useful model systems for studying cell differentiation as well as tissue development and function. Adipogenesis is mainly controlled by a cascade of sequentially expressed transcription factors (TFs) ^8^. Although many TFs have been implicated in adipogenesis, PPARγ and C/EBPα are primary drivers of the induction of thousands of adipocyte genes ^9,10^. Adipose tissues, including white adipose tissue (WAT) and brown adipose tissue (BAT), are dynamic endocrine organs that regulate thermogenesis, energy metabolism and homeostasis ^11^. The study of adipogenesis and adipose tissue *in vivo* requires the isolation of tissues of the adipose lineage at particular developmental and functional stages. Myogenic factor 5 (*Myf5*) promoter-driven Cre (*Myf5*-*Cre*) allows factors to be expressed or deleted in mouse preadipocytes, enabling a focus on adipogenesis. Conversely, *Adiponectin* promoter-driven Cre (*Adiponectin*-*Cre*) is generally expressed in differentiated adipocytes but not precursor cells, permitting study of adipocyte function ^12^. By crossing *Mll4* conditional knockout (KO) mice with *Myf5-Cre* or *Adiponectin*-*Cre* mice, we have shown that MLL4 is required for adipose tissue development but largely dispensable for adipose tissue maintenance ^4,5^. However, the roles of MLL3/4 enzymatic activities and MLL3/4-mediated H3K4me1 in adipose tissue development and function are unclear.

By tissue-specific ectopic expression of a histone H3.3 lysine-to-methionine mutant (H3.3K4M) in mice, we show that depletion of H3K4 methylation by H3.3K4M inhibits adipose tissue development. By tissue-specific deletion of the enzymatic SET domain of MLL3/4 in mice, we further show that the SET domain is required for adipose tissue development. Mechanistically, expression of H3.3K4M or deletion of the SET domain prevents MLL3/4-mediated enhancer activation in adipogenesis by destabilizing MLL3/4 proteins. Interestingly, H3.3K4M does not affect adipose tissue maintenance nor the thermogenic function of BAT.

## Results

### Histone H3.3K4M and H3.3K36M mutations impair adipogenesis

Previous studies reported that ectopic expression of histone H3.3 lysine-to-methionine (K-to-M) mutant specifically depletes endogenous lysine methylation in cells ^13, 14^. To understand the role of site-specific histone methylation in adipogenesis, we used retroviruses to stably express wild type (WT) or K-to-M mutant (K4M, K9M, K27M or K36M) of histone H3.3 in brown preadipocytes. The expression levels of FLAG-tagged H3.3 were much lower than that of endogenous H3 (Figure 1a). Consistent with previous reports ^13, 14^, ectopic expression of H3.3K4M selectively decreased global H3K4me1, H3K4me2, and H3K4me3 levels, while H3.3K9M and H3.3K27M selectively decreased global levels of H3K9me2 and H3K27me3, respectively. Ectopic expression of H3.3K36M selectively depleted global H3K36me2 and moderately increased H3K27me3 levels (Figure 1a and Supplementary Fig S1a). Cells were induced to undergo adipogenesis. As shown in Figure 1b-c, H3.3K4M-expressing (K4M) cells showed severe defects in adipogenesis and associated expression of adipogenesis markers *Pparg*, *Cebpa* and *Fabp4.* Consistent with our recent report ^29^, H3.3K36M-expressing cells also showed defects in adipogenesis. These results indicate that histone H3.3K4M and H3.3K36M mutations impair adipogenesis.

**Figure 1.**
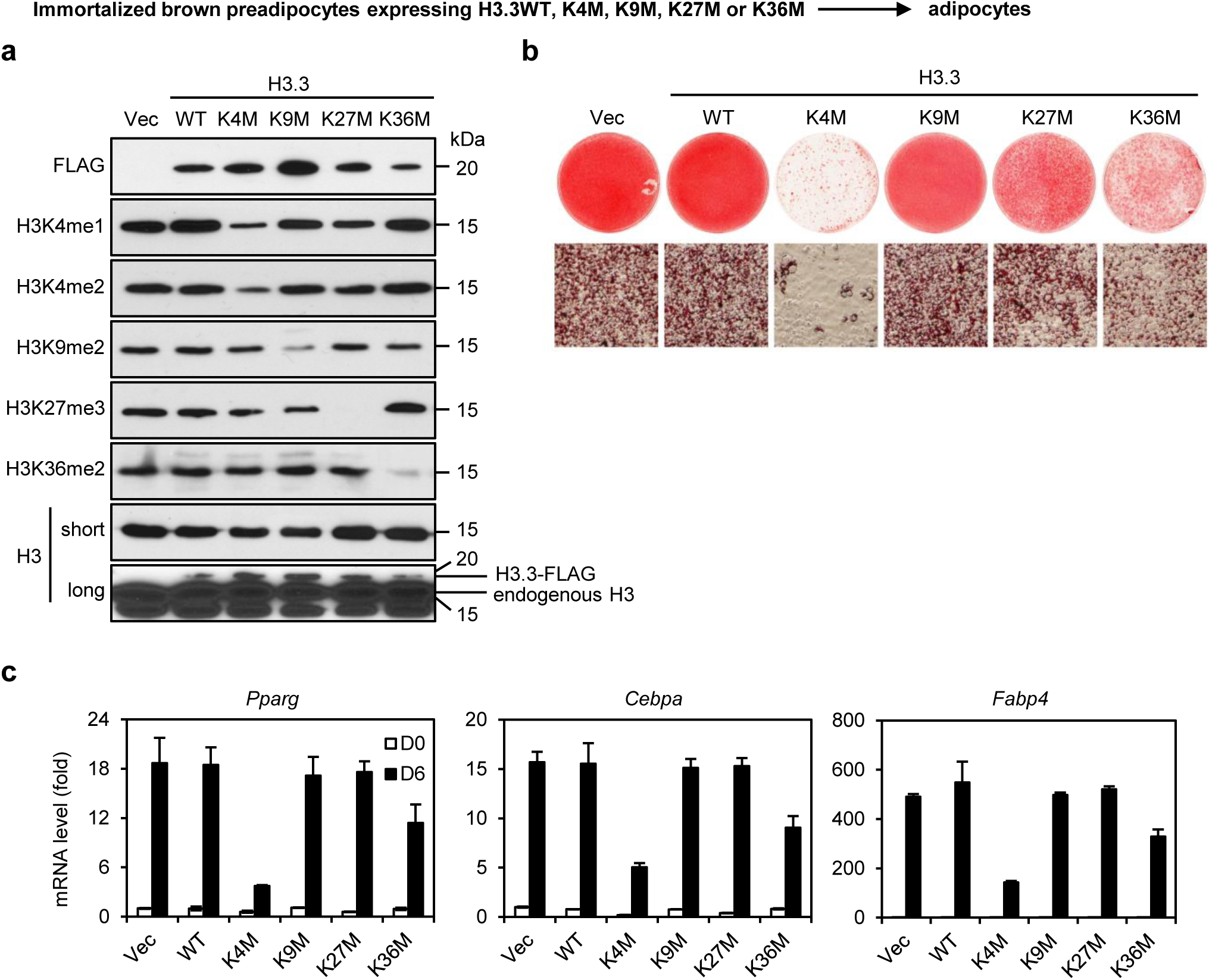
Histone H3.3K4M and H3.3K36M mutations impair adipogenesis. Immortalized brown preadipocytes were infected with retroviral vector (Vec) expressing FLAG-tagged wild type (WT) or K-to-M mutant histone H3.3, followed by adipogenesis assay. (a) Histone extracts from preadipocytes were subjected to Western blot analyses using antibodies indicated on the left. Long exposure of histone H3 Western blot reveals the relative levels of ectopic H3.3 and endogenous H3. (b) 6 days after induction of differentiation, cells were stained with Oil Red O. Upper panels, stained dishes; lower panels, representative fields under microscope. (c) qRT-PCR of *Pparg, Cebpa* and *Fabp4* expression at day 0 (D0) and day 6 (D6) of adipogenesis. Quantitative PCR data in all figures except Figure 8 are presented as means ± SD.

### H3.3K4M inhibits adipose tissue and muscle development

Next, we investigated whether H3.3K4M affects adipose tissue development *in vivo.* We generated a conditional H3.3K4M transgenic mouse line, lox-STOP-lox-H3.3K4M (LSL-K4M) for tissue-specific expression of H3.3K4M. The insertion of 4 copies of SV40 stop signals (STOP) flanked by two loxP sites prevents the CAG promoter-driven expression of FLAG-tagged H3.3K4M in the absence of Cre (Figure 2a-b). We crossed LSL-K4M mice with Myf5-Cre mice to induce H3.3K4M expression specifically in somitic precursor cells of brown adipose tissue (BAT) and skeletal muscle ^4^. No LSL-K4M;*Myf5*-*Cre* mice were observed at the weaning age. Newborn (P0) LSL-K4M;*Myf5-Cre* pups were obtained but died immediately after birth from breathing malfunctions due to deficient muscle groups in the rib cage (Figure 2c-d). Immunohistochemical analysis of cervical regions of E18.5 LSL-K4M;*Myf5*-*Cre* embryos revealed marked reduction in BAT and back muscle mass (Figure 2e-f), indicating that expressing H3.3K4M in progenitor cells prevents adipose tissue and muscle development *in vivo.*

**Figure 2.**
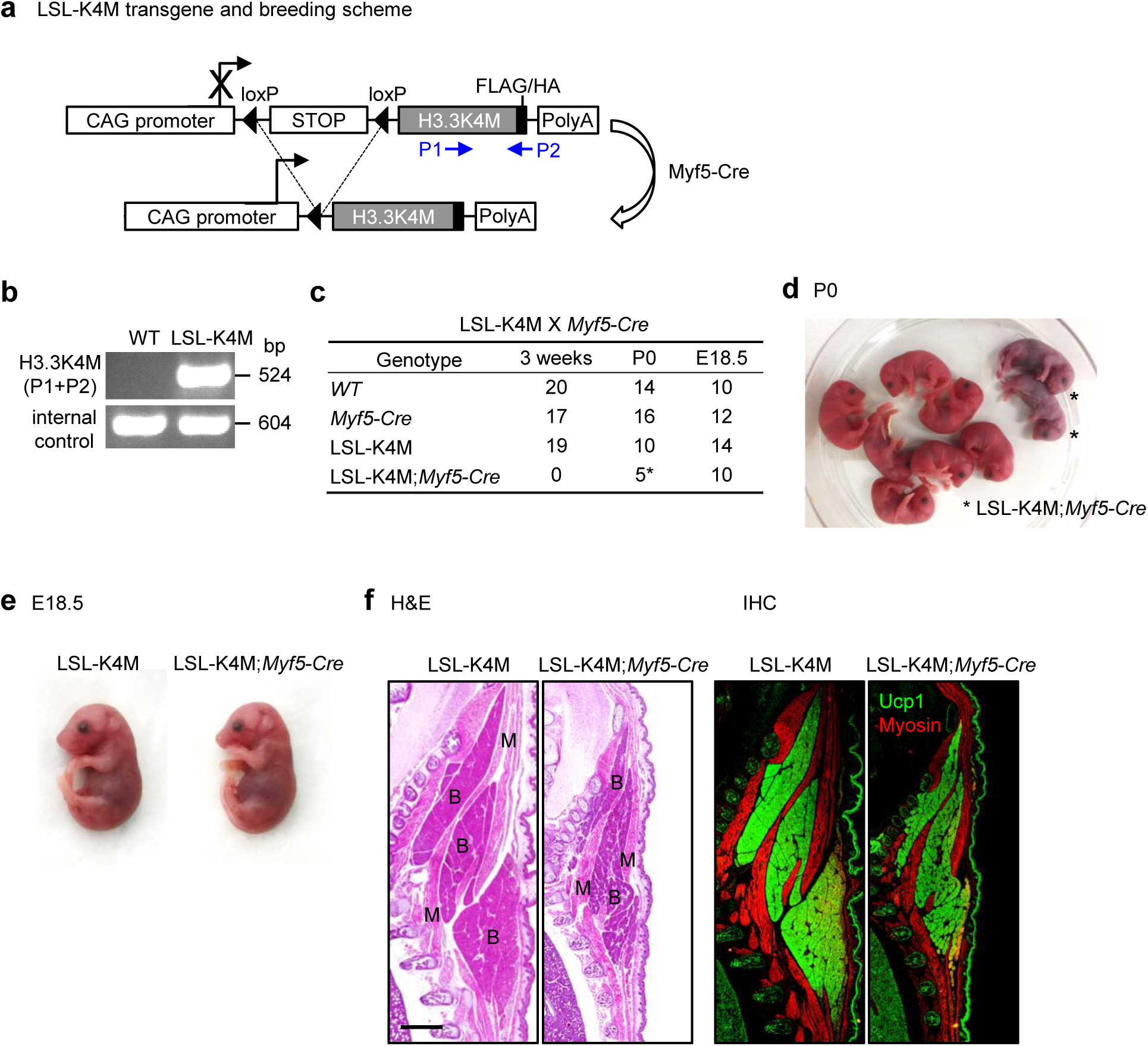
H3.3K4M prevents adipose tissue and muscle development. (a) Schematic of lox-STOP-lox-H3.3K4M (LSL-K4M) transgene and breeding scheme. The LSL-K4M transgene consists of the following elements from 5’ to 3’: a CAG promoter, quadruple copies of SV40 stop signals flanked by two loxP sites, H3.3K4M with C-terminal FLAG and HA tags, and polyadenylation signal. LSL-K4M transgenic mice were crossed with *Myf5*-*Cre* to generate mice expressing ectopic H3.3K4M in brown adipose tissue (BAT) and muscle. The locations of PCR genotyping primers P1 and P2 are indicated by arrows. (b) PCR genotyping of LSL-K4M transgenic mice. (c) Genotype of progeny from crossing between LSL-K4M and *Myf5*-*Cre* at 3 weeks age, new born pups (P0) and E18.5 embryos. LSL-K4M;*Myf5*-*Cre* mice died soon after birth from breathing malfunction due to defects in muscles of the rib cage. (d) Representative morphology of P0 pups. (e) Representative morphology of E18.5 embryos. (f) Histological analysis of E18.5 embryos. Sagittal sections of cervical/thoracic area were stained with H&E (left panels) or with antibodies against the BAT (B) marker UCP1 (green) and the muscle (M) marker Myosin (red) (right panels). *Scale bar* = 80 μm.

### H3.3K4M destabilizes MLL3/4 proteins in adipogenesis

To confirm that H3.3K4M prevents adipogenesis in a cell autonomous manner, we crossed LSL-K4M with *Cre*-*ER* mice to obtain primary LSL-K4M;*Cre*-*ER* brown preadipocytes. After immortalization, cells were treated with 4-hydroxytamoxifen (4OHT) to delete the STOP cassette and induce FLAG-tagged H3.3K4M expression. As expected, induction of H3.3K4M expression decreased global levels of endogenous H3K4me1/2/3. Interestingly, H3.3K4M expression also decreased H3K27ac levels (Figure 3a). Consistent with our previous findings observed in embryonic stem (ES) cells ^15^, expressing H3.3K4M destabilized endogenous MLL3/4 as well as the MLL3/4-associated protein UTX in cells. However, H3.3K4M did not affect protein levels of other members of the mammalian Set1-like H3K4 methyltransferase family, including SET1A, SET1B, and MLL1 ^16^ (Figure 3b).

**Figure 3.**
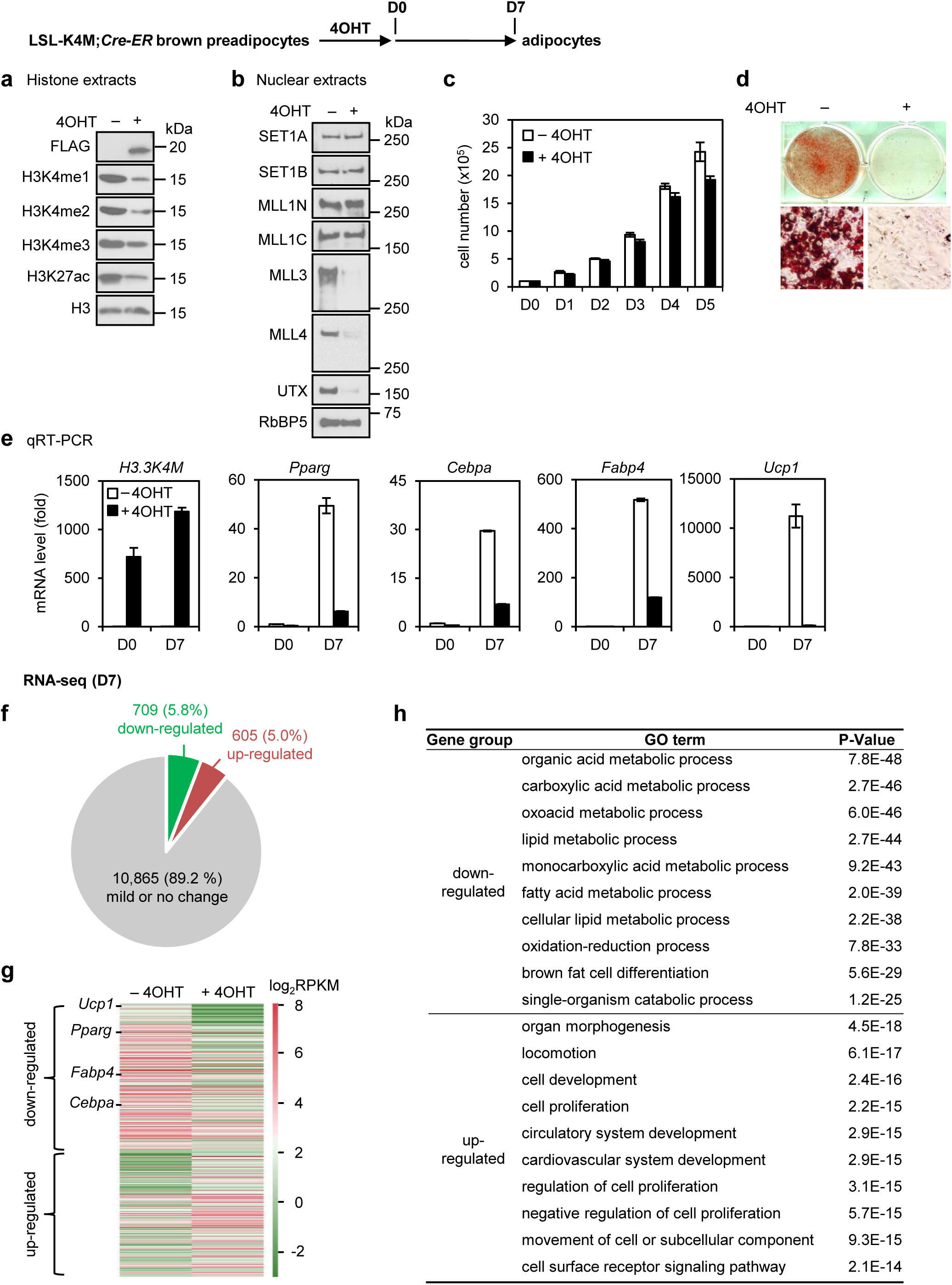
H3.3K4M destabilizes MLL3/4 proteins in adipogenesis. Immortalized LSL-K4M;*Cre*-*ER* brown preadipocytes were treated with 4-hydroxytamoxifen (4OHT) to induce ectopic H3.3K4M expression, followed by adipogenesis assay. (a–b) H3.3K4M destabilizes MLL3/4 proteins. Histone extracts (a) or nuclear extracts (b) were analyzed by Western blot using antibodies indicated on the left. (c–e) H3.3K4M prevents adipogenesis and induction of adipocyte genes. (c) Cell growth rates. 5 × 10^5^ preadipocytes were plated at D0 and the cumulative cell numbers were determined every day for 5 days. (d) Oil red O staining at day 7 (D7) of adipogenesis. (e) qRT-PCR of H3.3K4M-FLAG, *Pparg*, *Cebpa*, *Fabp4* and *Ucp1* expression at D0 and D7 of adipogenesis. (f-h) RNA-seq analyses were performed at D7 of adipogenesis. (f-g) Identification (f) and heat map (g) of down- or up-regulated genes in H3.3K4M-expressing cells. The cut-off for differential expression is 2.5- fold. (h) Gene ontology (GO) analysis of gene groups defined in (f).

Ectopic expression of H3.3K4M had little effect on cell proliferation (Figure 3c), but prevented adipogenesis and the induction of adipocyte marker genes such as *Pparg*, *Cebpa* and *Fabp4* as well as BAT-specific marker gene *Ucp1* (Figure 3d-e). We confirmed these H3.3K4M-driven adipogenesis defects using independent brown preadipocyte lines stably expressing WT H3.3 or H3.3K4M (Supplementary Fig S1). To investigate how H3.3K4M inhibits adipogenesis, we further performed RNA-seq analysis of LSL-K4M;*Cre*-*ER* preadipocytes treated with or without 4OHT. Using a 2.5-fold cut-off for differential gene expression from RNA-seq analysis, we defined genes up-regulated (605/5.0%) or down-regulated (709/5.8%) by H3.3K4M at D7 of differentiation (Figure 3f-g). Gene ontology (GO) analysis showed that down-regulated genes were strongly associated functionally with fat cell differentiation and lipid metabolism (Figure 3h). Because MLL3/4 are essential for adipogenesis ^4^, these data suggest that H3.3K4M inhibits adipogenesis at least in part by destabilizing MLL3/4 proteins.

### H3.3K4M prevents MLL3/4-mediated enhancer activation in adipogenesis

Next, we investigated whether H3.3K4M affects MLL3/4-mediated enhancer activation in adipogenesis. We performed ChIP-seq analyses of enhancer marks H3K4me1 and H3K27ac in LSL-K4M;*Cre*-*ER* cells treated with or without 4OHT at D4 of adipogenesis. Since ChIP-seq analysis did not consider the global differences between samples, we used histone Western blot data as a normalization control for more accurate quantitative analysis of H3K4me1 and H3K27ac (Supplementary Fig S2). By comparing with the published MLL4 ChIP-seq data during adipogenesis ^4,6^, we identified 6,686 MLL4^+^ active enhancers during adipogenesis. 4OHT-induced H3.3K4M expression prevented H3K4me1 and H3K27ac accumulation on MLL4^+^ active enhancers during adipogenesis (Figure 4a-b). Similar results were observed on *Pparg* and *Cebpa* loci (Figure 4c). Further, ChIP-qPCR analyses revealed that, on representative MLL4^+^ active enhancers (e1-e5) located on gene loci of master adipogenic regulators PPARγ and CEBPα ^7^, H3.3K4M markedly reduced the occupancy of MLL4, MLL3/MLL4-mediated H3K4me1, CBP/p300-mediated H3K27ac, BRD4, the MED1 subunit of the Mediator coactivator complex, and Pol II (Figure 4c-d). H3.3K4M also decreased eRNA production from MLL4^+^ adipogenic enhancers (Figure 4e). Together, these results suggest that H3.3K4M prevents MLL3/4-mediated enhancer activation in adipogenesis.

**Figure 4.**
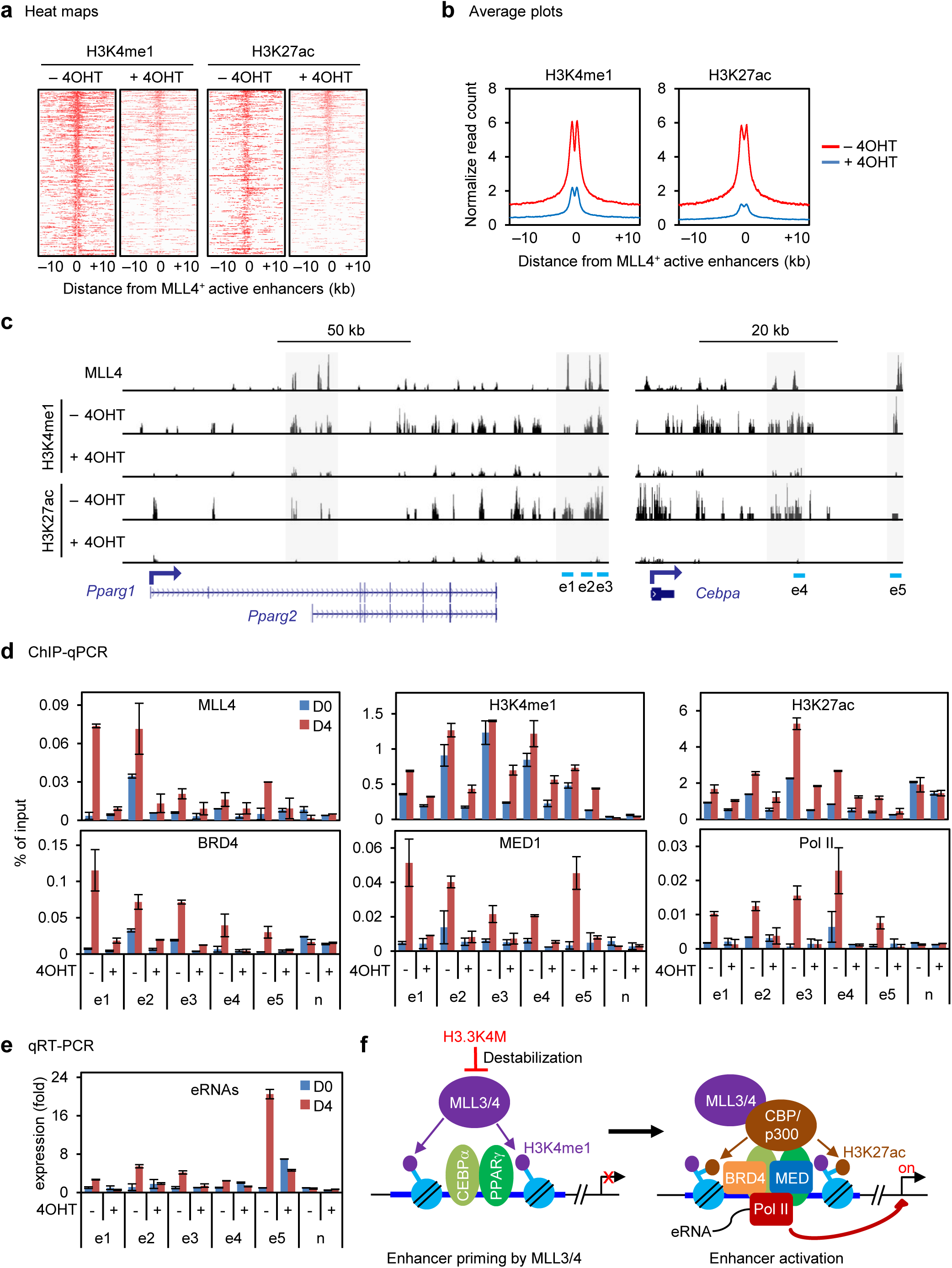
H3.3K4M prevents MLL3/4-mediated enhancer activation in adipogenesis. 4OHT-treated LSL-K4M;*Cre*-*ER* brown preadipocytes were collected at D4 of adipogenesis for ChIP-seq of H3K4me1 and H3K27ac (a-b), ChIP of MLL4, H3K4me1, H3K27ac, BRD4, MEDI and Pol II, and qRT-PCR of eRNAs (c-e). (a-b) Heat maps (a) and average profiles (b) around MLL4^+^ active enhancers during adipogenesis. (c) Genome browser view of H3K4me1 and H3K27ac on *Pparg* and *Cebpa* gene loci during adipogenesis with schematic of genomic locations of representative MLL4+ active enhancers (e1-e5). MLL4 binding data were obtained from ^4^. (d) ChIP-qPCR analyses of indicated factors are shown on enhancers e1-e5 at D0 and D4 of adipogenesis. An enhancer of constitutively expressed gene *Jak1* was chosen as negative control (n). (e) qRT-PCR of eRNA transcription on enhancers e1-e5 at D0 and D4 of adipogenesis. (f) Proposed model showing that H3.3K4M prevents enhancer activation in adipogenesis by destabilizing MLL3/4.

### Deletion of the enzymatic SET domain of MLL3/4 prevents adipose tissue and muscle development

MLL3 and MLL4 are partially redundant and are major H3K4 mono-methyltransferases on enhancers in cells ^4^. In a separate attempt to investigate the functional role of MLL3/4 enzymatic activity in differentiation and development *in vivo,* we used two conditional KO mouse lines targeting the enzymatic SET domain of MLL3 and MLL4 (*Mll3^f/f^* and *Mll4SET^f/f^,* Figure 5a-b). In the *Mll3^f/f^* mice, exons 57 and 58, which encode critical amino acids of the SET domain, were flanked by two loxP sites ^17^. In the *Mll4SET^f/f^* mice, exons 50 and 51, which encode the entire SET domain, were flanked by two loxP sites. Cre-mediated deletion of these exons would result in the production of enzyme-dead MLL3/4.

**Figure 5.**
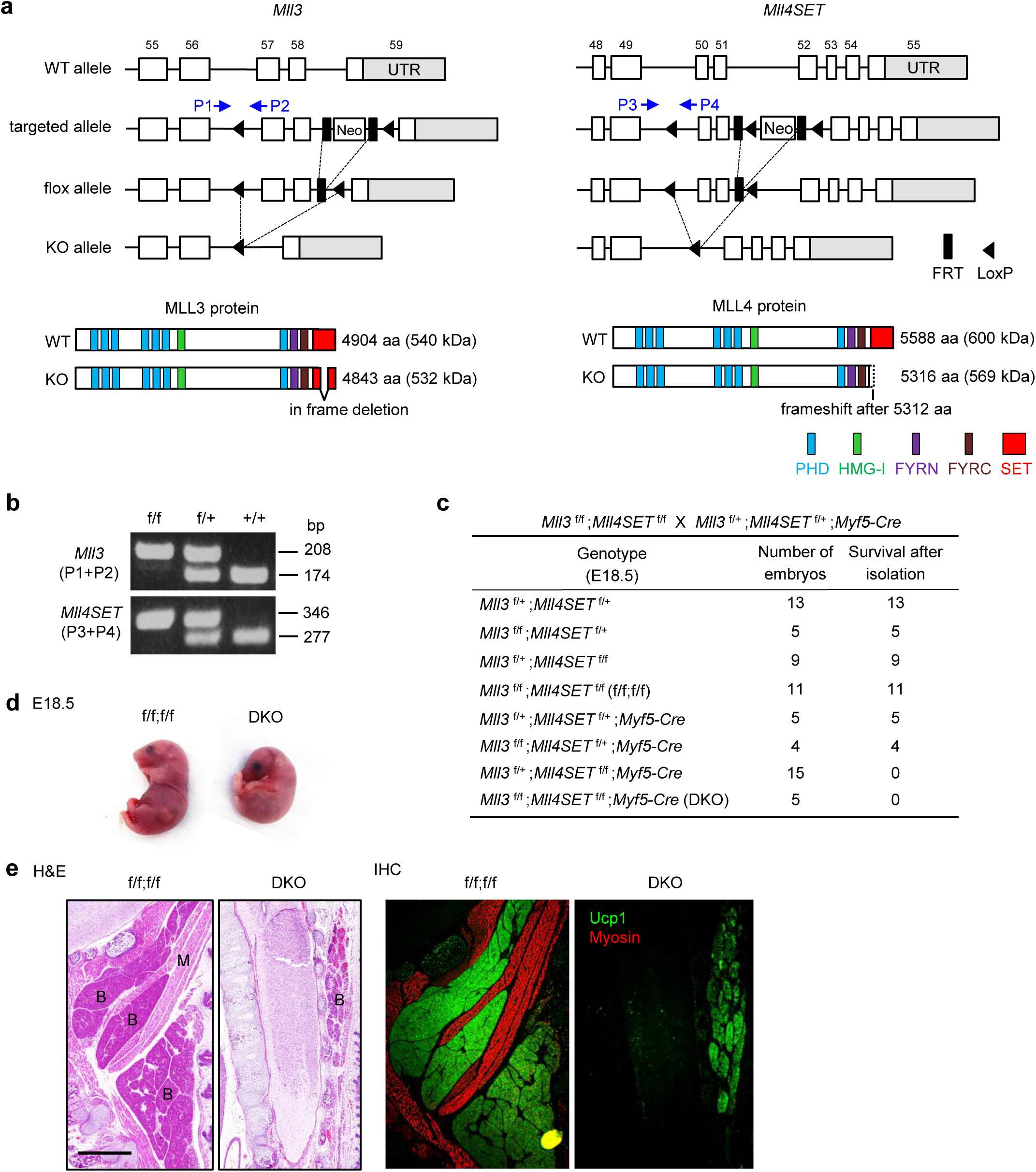
Deletion of the enzymatic SET domain of MLL3/4 prevents adipose tissue and muscle development. (a) Conditional KO mouse lines targeting the SET domain of MLL3 and MLL4. Schematics of WT allele, targeted allele, conditional KO (flox) allele and KO allele are shown in upper panels. Deletion of neomycin selection cassette by FLP recombinase generates the flox allele. The locations of PCR genotyping primers P1-P4 are indicated by arrows. Expected protein domains and molecular weights are shown in bottom panels. (b) PCR genotyping of flox and WT alleles using P1-P4 primers. (c) Genotypes of progeny at E18.5 from crossing between *Mll3^f/f^;Mll4SEf^f/f^* (f/f;f/f) and *Mll3*^*f*/+^;*Mll4SEf*^*f*/+^;*Myf5*-*Cre*. *Mll3*^*f*/+^;*Mll4SET^f/f^;Myf5*-*Cre* and *Mll3^f/f^;Mll4SET^f/f^;Myf5*-*Cre* (conditional double KO, DKO) mice died immediately after cesarean section from breathing malfunction due to defects in muscles of the rib cage. (d) Representative morphology of E18.5 embryos. (e) Histological analysis of E18.5 embryos. Sagittal sections of cervical/thoracic area were stained with H&E (upper panels) or with antibodies against the BAT (B) marker UCP1 (green) and the muscle (M) marker Myosin (red) (lower panels). *Scale bar* = 80 μm.

We first crossed *Mll4SEf*^f/f^ with *Myf5*-*Cre* mice. The resulting *Mll4SET*^f/f^;*Myf5*-*Cre* mice survived until birth. E17.5~18.5 *Mll4SET*^f/f^;*Myf5*-*Cre* embryos were unable to breathe and died immediately after isolation, displaying an abnormal hunched posture and severe reduction of back muscles (Supplementary Fig S3a-c). These embryos showed only a moderate decrease of BAT mass compared to WT, possibly due to a compensatory effect of MLL3 (Supplementary Fig S3c). To eliminate the compensatory effect, we crossed *Mll4SET*^f/f^ with *Mll3*^f/f^ and *Myf5*-*Cre* mice to delete both *Mll3* and *Mll4* genes in progenitor cells of BAT and muscle lineages. The resulting E18.5 *Mll3^f/f^;Mll4SEf*^f/f^;*Myf5*-*Cre* (conditional double KO, DKO) embryos were unable to breathe and died immediately after isolation. These embryos showed profound reduction of BAT as well as muscle mass (Figure 5c-e). These data indicate that deletion of the enzymatic SET domain of MLL3/4 prevents adipose tissue and muscle development.

### Deletion of the enzymatic SET domain inhibits adipogenesis by destabilizing MLL3/4

To confirm that deletion of the enzymatic SET domain of MLL3/4 prevents adipogenesis in a cell autonomous manner, we crossed *Mll3*^f/f^;*Mll4SET*^f/f^ mice with *Cre*-*ER* mice to obtain primary *Mll3*^f/f^;*Mll4SET*^f/f^;*Cre*-*ER* brown preadipocytes. After immortalization, cells were treated with 4OHT to delete exons encoding the SET domain of MLL3/4. Consistent with our previous finding that MLL3 and MLL4 are major H3K4 mono- and di-methyltransferases in cells ^4,5^, deletion of the enzymatic SET domain in preadipocytes decreased global levels of endogenous H3K4me1/2 but not H3K4me3 (Figure 6a). In addition, deleting the enzymatic SET domain destabilized endogenous MLL3/4 and reduced protein levels of MLL3/4-associated UTX in cells (Figure 6b), which is consistent with a recent report that deleting the SET domain destabilizes MLL3/4 in ES cells ^18^.

**Figure 6.**
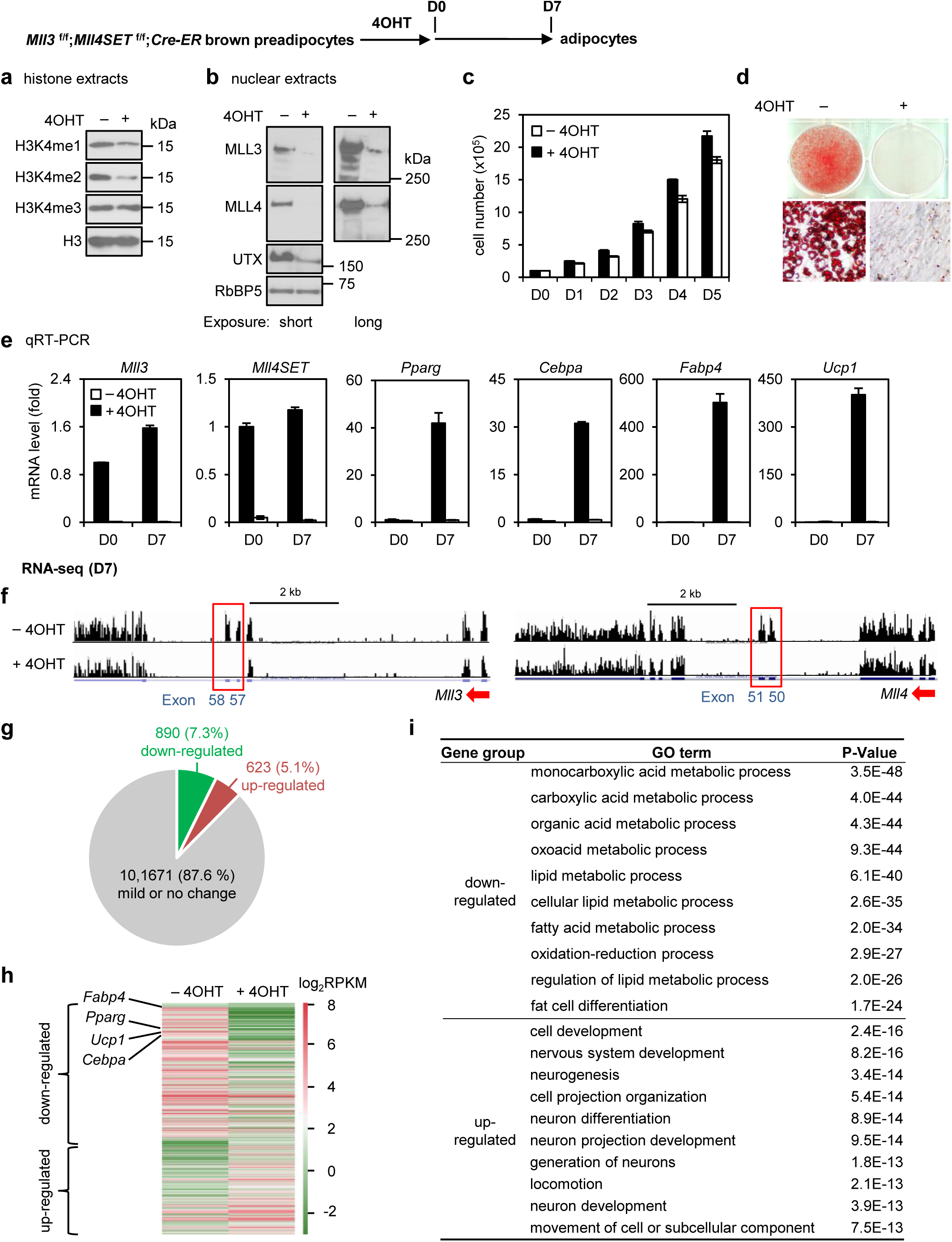
Deletion of the enzymatic SET domain inhibits adipogenesis by destabilizing MLL3/4. Immortalized *Mll3^f/f^;Mll4SET^f/f^;Cre*-*ER* brown preadipocytes were treated with 4OHT to induce deletion of the exons encoding the SET domain of MLL3 and MLL4 proteins, followed by adipogenesis assay. (a–b) Deletion of the enzymatic SET domain destabilizes MLL3/4 proteins. Histone extracts (a) or nuclear extracts (b) from preadipocytes were analyzed by Western blot using antibodies indicated on the left. (c-e) Deletion of the SET domain of MLL3/4 prevents adipogenesis. (c) Cell growth rates. (d) Oil red O staining at D7 of adipogenesis. (e) qRT-PCR of *Mll3*, *Mll4SET*, *Pparg*, *Cebpa*, *Fabp4* and *Ucp1* expression at D0 and D7 of adipogenesis. (f-i) RNA-seq analyses were performed at D7 of adipogenesis. (f) Genome browser views of RNA-Seq analysis on *Mll3* and *Mll4* loci. The targeted exons are highlighted in red boxes. (g-h) Schematic of identification (f) and heat map (h) of down- or up-regulated genes in double KO (DKO) cells. The cut-off for differential expression is 2.5-fold. (i) GO analysis of gene groups defined in (g).

Deletion of the enzymatic SET domain of MLL3/4 had little effect on cell proliferation (Figure 6c), but prevented adipogenesis and the induction of *Pparg*, *Cebpa*, *Fabp4*, and *Ucp1* (Figure 6d-e). RNA-seq analysis at D7 of differentiation confirmed the deletion of target exons of both *Mll3* and *Mll4* loci (Figure 6f). We further performed RNA-seq analysis of *Mll3^f/f^;Mll4SEf^f/f^;Cre*-*ER* preadipocytes treated with or without 4OHT. Using a 2.5-fold cut-off for differential expression from RNA-seq analysis, we defined genes up-regulated (623/5.1%) or down-regulated (890/7.3%) by deletion of the enzymatic SET domain at D7 of differentiation (Figure 6g-h). GO analysis showed that down-regulated genes were strongly functionally associated with fat cell differentiation and lipid metabolism (Figure 6i). These data suggest that deletion of the enzymatic SET domain inhibits adipogenesis by destabilizing MLL3/4.

### H3.3K4M expression mimics MLL3/4 SET domain deletion in preventing adipogenesis

Next, we generated scatter plots of gene expression changes using RNA-seq data from cells with deletion of MLL3/4 SET domain (DKO) and cells with expression of H3.3K4M (K4M) (Figure 7a). A combined scatter plot revealed that down- or up-regulated genes were highly correlated between DKO and K4M cells (Figure 7b). We found that the majority of down or up-regulated genes at D7 of differentiation were shared by DKO and K4M (Figure 7c-d). The shared down-regulated genes were highly associated functionally with fat cell differentiation and lipid metabolism whereas shared up-regulated genes were highly associated with cell proliferation and development (Figure 7e-f). These data suggest that ectopic H3.3K4M mimics the effect of Mll3/4 SET domain deletion in preventing adipogenesis.

**Figure 7.**
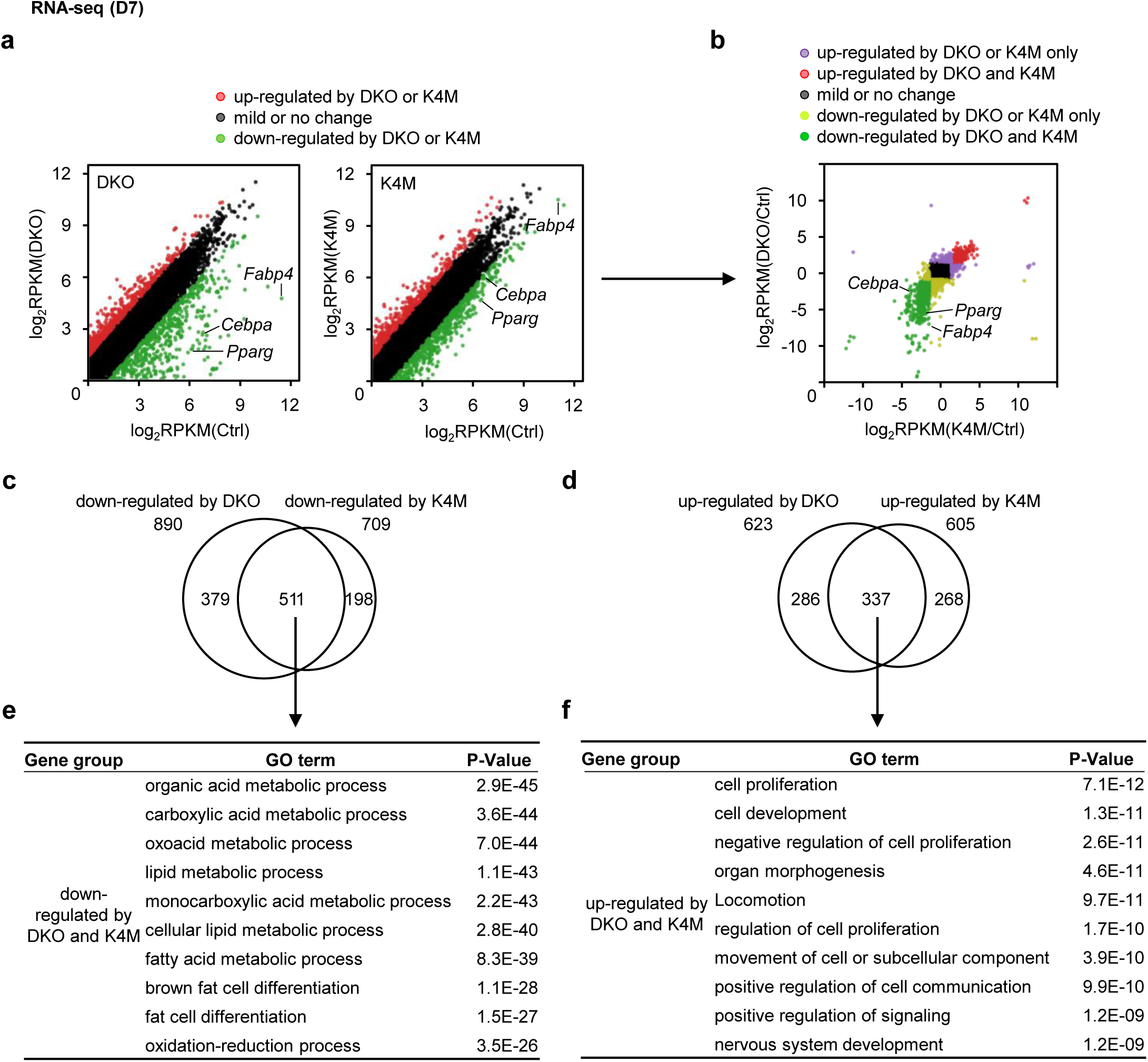
H3.3K4M expression mimics MLL3/4 SET domain deletion in preventing adipogenesis. (a) Scatter plots of down- or up-regulated genes in H3.3K4M-expressing or MLL3/4 SET domain DKO cells compared to the respective control (Ctrl) cells. RNA-seq data were from Figures 3 and 6. The cut-off for differential expression is 2.5-fold. Down- or up-regulated genes are depicted in green and red, respectively. (b) Combined scatter plot showing correlation of fold changes of gene expression by H3.3K4M expression and by MLL3/4 SET domain DKO. (c-d) Venn diagram showing the overlap of down- or up-regulated genes in H3.3K4M-expressing and MLL3/4 SET domain DKO cells. (e-f) GO analysis of gene groups defined in (c-d).

### H3.3K4M does not affect adipose tissue maintenance and function

We also investigated the role of H3K4 methylation in adipose tissue maintenance and function in mice. For this purpose, we crossed LSL-K4M mice with *Adipoq*-*Cre* mice to achieve adipocyte-selective expression of H3.3K4M *in vivo.* Histone Western blot of BAT from *LSL*-*K4M;Adipoq*-*Cre* mice showed that ectopic expression of H3.3K4M depleted endogenous H3K4me1/2/3 and MLL3/4 levels in BAT (Figure 8a-b). LSL-K4M;*Adipoq*-*Cre* mice did not show any discernable differences in body weight, fat/lean mass, or adipose tissue mass relative to control (LSL-K4M) mice at 8-10 weeks of age (Figure 8c-e). In the BAT, induction of H3.3K4M expression was successful in *LSL*-*K4M;Adipoq*-*Cre* mice, but the expression levels of adipocyte identity genes *Pparg*, *Cebpa*, and *Fabp4* were similar between *LSL*-*K4M;Adipoq*-*Cre* and control mice (Figure 8f). This phenotype is consistent with what we have reported for the *Mll4SET^f/f^;.Adipoq*-*Cre* mice, in which MLL4 is dispensable for the maintenance of differentiated adipocytes *in vivo* ^5^.

**Figure 8.**
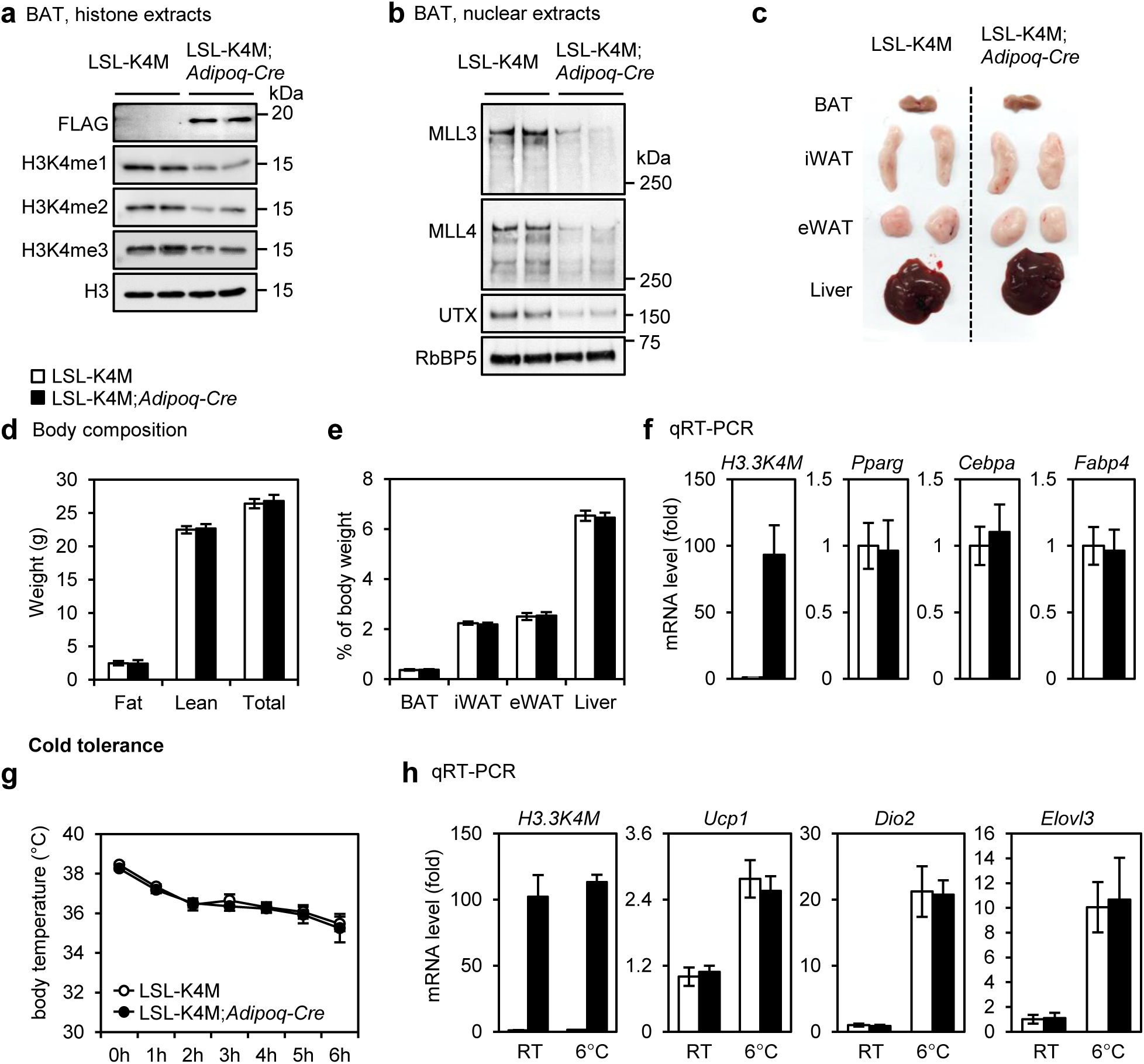
H3.3K4M does not affect adipose tissue maintenance and function. All data were from 8 to 10-week-old male mice fed with regular diet. (a-b) Histone extracts (a) or nuclear extracts (b) from BAT were analyzed by Western blot using antibodies indicated on the left. (c) Representative picture of BAT, inguinal WAT (iWAT), epididymal WAT (eWAT), and liver. (d) Fat mass, lean mass, and total body weight were measured by MRI (*n*=9 per group). (e) The average tissue weights are presented as % of body weight (*n*=6 per group). (f) qRT-PCR of *H3.3K4M*, *Pparg*, *Cebpa*, and *Fabp4* expression in BAT (*n*=6 per group). (g-h) Cold tolerance test. Mice were housed at room temperature (RT, 22°C) and then in a cold room (6°C) for 6h (*n*=6 per group). (g) Body temperatures. (h) qRT-PCR of gene expression in BAT after 6h. All values in Figure 8 are presented as mean ± S.E.M.

We further asked about the functional consequence of ectopic H3.3K4M expression in adipocytes *in vivo.* For this purpose, we acutely exposed the mice to environmental cold (6°C) up to 6 h. LSL-*K4M;Adipoq*-*Cre* mice maintained normal body temperatures, were cold tolerant and behaved similarly as control mice in the cold tolerance test (Figure 8g). After cold exposure, expression levels of thermogenesis genes *Ucp1*, *Dio2*, and *Elovl3* were similarly induced in BAT of LSL-K4M;*Adipoq*-*Cre* and control mice (Figure 8h). Together, our data indicate that while H3.3K4M prevents adipose tissue development, it does not affect the maintenance and function of adipose tissues. Our data also suggest that H3K4 methylation is dispensable for the maintenance and function of differentiated adipocytes.

## Discussion

Site-specific histone methylations are generally correlated with gene activation or gene repression. To investigate the role of site-specific histone methylations in cell differentiation and development, we screened several K-to-M mutants of H3.3 and found that H3.3K4M and H3.3K36M mutations impair adipogenesis in cell culture. Mechanistically, H3.3K4M destabilizes MLL3/4 proteins but not other members of the mammalian Set1-like H3K4 methyltransferase family, including SET1A, SET1B, and MLL1. Consequently, H3.3K4M prevents MLL3/4-mediated enhancer activation in adipogenesis. Using tissue-specific expression of H3.3K4M in mice, we next showed that H3.3K4M inhibits adipose tissue and muscle development. However, H3.3K4M does not affect adipose tissue maintenance nor the thermogenic function of BAT. Using tissue-specific deletion of the enzymatic SET domains of MLL3/4 in mice, we demonstrated that the SET domains are required for adipose tissue and muscle development. Mechanistically, deletion of the SET domains destabilizes MLL3/4 proteins. Notably, H3.3K4M expression mimics MLL3/4 SET domain deletion in preventing adipogenesis. Together, our findings suggest that H3.3K4M destabilizes enhancer epigenomic writers MLL3/4 and impairs adipose tissue development (Figure 4f).

Consistent with our observation in preadipocytes, it was shown recently that MLL3/4 proteins with SET domain deletion are unstable in ES cells ^18^. However, another recent report contradictorily showed no MLL3/4 protein stability defects in ES cells with MLL3/4 SET domain deletion ^19^. To find out what caused such discrepancies, we examined the different designs for SET domain deletion on MLL3/4 proteins in these three studies. We showed that deleting exons 57 and 58 results in a destabilized MLL3 protein in cells (Figure 6b). In contrast, Rickels et al. deleted exons 56, 57 and 58 without affecting MLL3 protein stability in cells ^19^. The full-length mouse MLL4 protein has 5588aa. Dorighi et al. showed that a truncated MLL4 protein containing amino acids 1-5482 is unstable in ES cells ^18^. In contrast, Rickels et al. showed that a more severely truncated MLL4 protein containing amino acids 1-5402 is stable in ES cells ^19^. Future studies are needed to verify whether the SET domain is required for MLL3/4 protein stability in cells. Consistent with our previous report that ectopic expression of H3.3K4M reduces endogenous levels of MLL3/4 and MLL3/4-associated UTX protein in ES cells ^15^, we now show that H3.3K4M destabilizes MLL3/4 and UTX proteins in preadipocytes and BAT. However, it remains to be determined mechanistically how H3.3K4M and the SET domains regulate MLL3/4 protein stability.

Although H3K4me1 is the predominant mark of a primed enhancer state, it is unclear whether H3K4me1 affects or simply correlates with enhancer activation in cell differentiation and development. Interestingly, the latest two studies from one group reported that MLL3/4-dependent H3K4me1 has an active role at enhancers by facilitating binding of the chromatin remodeler SWI/SNF complex and recruitment of the chromatin organization regulator cohesin complex to orchestrate long-range chromatin interactions in ES cells ^20,21^. These findings imply that H3K4me1 is not simply a correlative outcome in enhancer regulation. Conversely, using CRISPR to generate catalytically inactive MLL3/4, another recent study surprisingly reported that MLL3/4 proteins, rather than MLL3/4-mediated H3K4me1, are required for enhancer activation and gene transcription in undifferentiated ES cells. However, the role of MLL3/4-mediated H3K4me1 in ES cell differentiation was not elucidated ^18^. Thus, despite various efforts to uncover the role of H3K4me1 in enhancer function, it is still unclear whether H3K4me1 controls enhancer activation in cell differentiation. Therefore, future work will be needed to clarify the role of MLL3/4-mediated H3K4me1 in enhancer function during cellular differentiation and animal development.

## Methods

### Plasmids, Antibodies and Chemicals

The retroviral pQCXIP plasmids expressing FLAG-tagged wild type (WT) or mutants of histone H3.3 including K4M, K9M, K27M and K36 were described previously ^14^. The following homemade antibodies have been described: anti-MLL4#3 ^22^, anti-MLL3#3 ^16^ and anti-UTX ^23^. Anti-RbBP5 (A300-109A), anti-BRD4 (A301-985A100) and anti-MED1 (A300-793A) were from Bethyl Laboratories. Anti-SET1A/B antibodies were described previously ^24^. Anti-H3 (ab1791), anti-H3K4me1 (ab8895), anti-H3K4me2 (ab7766), anti-H3K27ac (ab4729) and H3K36me3 (ab9050) were from Abcam. Anti-MLL1N (A700-010), anti-MLL1C (A300-374A), anti-Pol II (17-672), anti-H3K4me3 (07-473), anti-H3K9me2 (17-648), anti-H3K27me3 (07-449) and anti-H3K36me2 (07-369) were from Millipore. Anti-FLAG-M2 (F3165) and (Z)-4-Hydroxytamoxifen (4OHT) (H7904) were from Sigma.

### Generation of Mouse Strains

To generate LSL-K4M transgenic mice, H3.3K4M and a 3’ FLAG tag (H3.3K4M-FLAG) were fused downstream of CAG promoter with a loxP-STOP-loxP cassette in the middle of the pBT346.6 plasmid (AST-3029, Applied StemCell) (Figure 2a). H3.3K4M-FLAG of pQCXIP-H3.3K4M was subcloned into the pBT346.6; after confirmation by DNA sequencing, the plasmid was linearized by SpeI and ScaI, gel purified and injected into zygotes harvested from C57BL/6 mice. Founder mice were identified by genotyping. For genotyping the *LSL*-*K4M* alleles, PCR was done using the following primers: 5’-CTAGCTGCAGCTCGAGTGAACCATGGC-3’ and 5’-TTCGCGGCCGCGAATTCCTAGGCGTAGTCG-3’. PCR amplified 524 bp from the *LSL-K4M* alleles.

*Mll3*^f/f^ mice were obtained from Jae W. Lee ^17^ (Figure 5a, left panel). To generate *Mll4SET* conditional KO mice, the loxP/FRT-flanked neomycin cassette was inserted at the 3’ end of exon 51 and the single loxP site was inserted at the 5’ end of exon 50 (Figure 5a, right panel). We electroporated the linearized targeted construct, which includes exons 50–51, into WT ES cells. After selection with G418, surviving clones were expanded for PCR genotyping to identify *Mll4SET*^floxneo/+^ ES cells, which were further micro-injected into mouse blastocysts following standard procedures. Mice bearing germline transmission (*Mll4SET*^floxneo/+^) were crossed with FLP1 mice (Jackson no. 003946) to generate *Mll4SEf*^f/+^ mice. For genotyping the *Mll3* and *Mll4SET* alleles, PCRs were done using the following primers: *Mll3* (5’-GTCATCGGTGTGGTCTGAATGA-3’ and 5’-AACCGGAAGGAGAAGCTTTATGA-3’) and *Mll4SET* (5’-CAGTTGAGCTAGTCAAGTGATT-3’ and 5’-TTCAATGTGGAGGGGAGTGACAG-3’). PCR amplified 174 bp from the wild-type *Mll3* and 208 bp from the *Mll3* floxed allele, or 277 bp from the wild-type *Mll4* and 346 bp from the *Mll4SET* floxed allele.

LSL-K4M mice and *Mll3*^f/f^;*Mll4SEf*^f/f^ mice were crossed with *Myf5*-*Cre* (Jackson no. 007893), *Cre*-*ER* (Jackson no. 008463), or *Adipoq*-*Cre* (Jackson no. 028020) to generate LSL-K4M;*Myf5*-*Cre*, LSL-K4M; *Cre*-*ER*, LSL-K4M;.*Adipoq*-*Cre*, *Mll4SEf*^f/f^;*Myf5*-*Cre*, *Mll3*^f/f^;*Mll4SEf*^f/f^;*Myf5*-*Cre*, or *Mll3*^f/f^;*Mll4SET*^f/f^;*Cre*-*ER* mice.

### Histology and Immunohistochemistry

E18.5 embryos were isolated and fixed in 4% paraformaldehyde, dehydrated in ethanol and embedded in paraffin for sectioning. Paraffin sections were stained with routine H&E or subjected to immunohistochemistry using anti-Ucp1 (ab10983; Abcam) and anti-Myosin (MF20; Developmental Studies Hybridoma Bank) antibodies as described ^4^.

### Immortalization of Primary Brown Preadipocytes and Adipogenesis

Primary brown preadipocytes were isolated from interscapular BAT of newborn LSL-K4M;*Cre*-*ER* or *Mll3*^f/f^;*Mll4SET*^f/f^;*Cre*-*ER* pups, and immortalized by SV40T. Isolation, immortalization and adipogenesis assays of brown preadipocytes were done as described ^25,26^.

### Western Blot and qRT-PCR

Western blot of histone modifications using acid extracts or of nuclear proteins using nuclear extracts were done as described ^15^. Total RNA was extracted using TRIzol (Invitrogen) and reverse transcribed using ProtoScript II first-strand cDNA synthesis kit (NEB), following the manufacturers’ instructions. qRT-PCR was done using the following SYBR green primers: *H3.3K4M* (forward, 5’-AACCTGTGTGCCATCCACG-3’, and reverse, 5’-CGACTTGTCATCGTCGTCCTT-3’), *Mll3* (forward, 5’-GATTGACGCCACACTCACAG-3’, and reverse, 5’-TTTCTGTATCCTCCGGTTGG-3’), and *Mll4SET* (forward, 5’-GGGTGGAGAGCTGTCAGAATTATT-3’, and reverse, 5’-CATGAGCGGTAACTCCATCAGA-3’). SYBR green primers for other genes were described previously ^27^.

### RNA-Seq and ChIP-seq

RNA-seq and ChIP-seq were performed as described in detail previously with the use of Illumina HiSeq 2500 ^4, 6, 7^. For RNA-seq, we purified mRNAs using Dynabeads mRNA purification kit (Invitrogen) then synthesized double-stranded cDNAs using SuperScript Double-stranded cDNA Synthesis kit (Invitrogen), following the manufacturers’ instructions. For ChIP-seq, we collected ChIP-DNA using Dynabeads Protein A (Invitrogen) then purified DNA using QIAquick PCR Purification Kit (QIAGEN), following the manufacturers’ instructions. Library construction for RNA-seq and ChIP-seq was completed using NEBNext Ultra II DNA Library Prep Kit (NEB), following the manufacturers’ instructions.

### Computational Analysis and Data Availability

For differentially expressed genes from RNA-seq data, gene ontology (GO) analysis was carried out using DAVID (https://david.ncifcrf.gov). To identify H3K4me1 or H3K27ac enriched regions at D4 of adipogenesis in LSL-K4M;CreER brown preadipocytes, we used ‘SICER’ method ^28^ with window size of 200 bp and with an estimated false discovery rate (FDR) threshold of 10^−3^. MLL4 ChIP-Seq data at D2 of adipogenesis was downloaded (GSE74189) ^4^ and the window size was chosen to be 50 bp. To define MLL4^+^ active enhancers during adipogenesis, we compared 13,871 MLL4 binding sites at D2 with 64,757 active enhancers (H3K4me1^+^H3K27ac^+^ at D4). 6,686 MLL4^+^ (10.3%, 6,686/64,757) sites were located on active enhancers. Heat maps were generated with 50 bp resolution and ranked according to the intensity of 6,686 MLL4^+^ active enhancers at the center. Average profiles were plotted using the number of ChIP-Seq reads from the center of 6,686 MLL4^+^ active enhancers to 10 kb on both sides. H3K4me1 and H3K27ac ChIP-seq signal intensity was normalized by histone Western blot for baseline modification levels to achieve more accurate quantitative analysis.

All datasets described in the paper have been deposited in NCBI Gene Expression Omnibus under accession number GSE110972.

## Acknowledgements

We thank Victoria Noe-Kim for helping computational analyses. This work was supported by the Intramural Research Program of NIDDK, NIH to KG.

## Author contributions

YJ and KG conceived and designed the experiments. Y.J., C.W., Y-K.P., L.Z., J-E.L., A.B. and, E.F. performed experiments. Y.J., C.W., Y-K.P., L.Z., J-E.L. and, K.G. analyzed the data. Y.J., and C.L. generated LSL-K4M mice and C.W., and C.L. generated *Mll4SET*^f/f^ mice. Y.J. performed computational analyses. Y.J., A.B., and K.G. wrote the manuscript. K.G. supervised all the experiments.

## Additional information

### Supplementary Information

